# Pasa: Leverage population pangenome graph to scaffold prokaryote genome assemblies

**DOI:** 10.1101/2023.07.09.548288

**Authors:** Van Hoan Do, Son Hoang Nguyen, Duc Quang Le, Tam Thi Nguyen, Canh Hao Nguyen, Tho Huu Ho, Vo Sy Nam, Trang Nguyen, Hoang Anh Nguyen, Minh Duc Cao

## Abstract

Whole genome sequencing has increasingly become the essential method for studying the genetic mechanisms of antimicrobial resistance and for surveillance of drug-resistant bacterial pathogens. The majority of bacterial genomes sequenced to date have been sequenced with Illumina sequencing technology, owing to its high-throughput, excellent sequence accuracy, and low cost. However, because of the short-read nature of the technology, these assemblies are fragmented into large numbers of contigs, hindering the obtaining of full information of the genome. We develop Pasa, a graph-based algorithm that utilizes the pangenome graph and the assembly graph information to improve scaffolding quality. By leveraging the population information of the bacteria species, Pasa is able to utilize the linkage information of the gene families of the species to resolve the contig graph of the assembly. We show that our method outperforms the current state of the art in terms of accuracy, and at the same time, is computationally efficient to be applied to a large number of existing draft assemblies.

## 1. Introduction

The increasing availability of DNA sequencing technologies has profoundly transformed biomedical research and in particular microbial genomics. The ability to decode the whole genomes of a large number of bacterial isolates enables the study of the genetic mechanisms of antimicrobial resistance in drug-resistant pathogens [1, 2], which have been rapidly emerging and are considered to be one of the main threats to public health in the next decades. Whole genome sequencing is also an effective tool for the surveillance of infectious diseases, and direct infection control measures in clinics [3–5]. Substantial global efforts have resulted in large amounts of sequencing data for genome assemblies being generated to provide insights into the resistant causes and effects. To date, short-read sequencing technology using Illumina platforms remains the most common method for whole genome sequencing owing to its high throughput, sequence accuracy, and cost-effectiveness. However, the relatively short read length cannot unambiguously resolve the repetitive sequences that are frequently present in most genomes. As a result, most genome assemblies are fragmented into large numbers of contigs and require additional information to improve their contiguity and completeness.

Much recent research has resulted in both technological and computational methodologies to generate better complete genome assemblies. Third-generation sequencing technologies such as Oxford Nanopore Technology and PacBio provide long reads spanning repetitive genomic regions. These long reads can be used for *de novo* assembling complete prokaryote genomes by long-read assemblers [6, 7], or for scaffolding fragmented assemblies from short-read sequencing [8–10]. However, higher costs and less mature ecosystems make long-read sequencing less attractive for large-scale sequencing projects. Long-range sequencing technologies such as mate-pairs, Hi-C [11], 10X Genomics linked-reads [12], and optical mapping [13] capture linkage information for inferring the distance and orientation between contigs in scaffolding assemblies. These technologies however often provide low-resolution information while having specific errors and biases, providing challenges to the computational analysis [14, 15], in addition to extra steps in data generation.

Computational approaches are also developed to utilize available public data resources as the reference to guide the scaffolding process. The rationale is to use an available, more complete assembly of the closest-possible genome as the backbone to place the contigs in the correct order. For *de novo* assembly, identifying a close-related genome can be a difficult task. Even if possible, the issue of reference bias can become severe for those that are highly structurally variable, such as in multi-drug resistance strains. To alleviate this problem, a class of scaffolding methods that use multiple references for scaffolding have been developed. The prominent methods suitable for microbial genomes include Ragout [16, 17] and multi-CSAR [18].

The idea of using the genome variant graph as a comprehensive but compact reference model to alleviate the bias issue of using individual references has been proposed in various applications of human genomics [19, 20] and microbial genomics [21, 22]. This technique has shown advantages over the traditional linear reference in *e*.*g*., alignment, variant calling and typing analyses. However, due to costly computation, the use of the variant graph usually requires high-performance computing resources and extensive running time. Instead, a pangenome graph at the gene level is adequate for almost all bacterial genomics analyses. It can provide the graph structure regarding the proximity relationships of all genes involved. A pangenome graph can be built from a set of bacterial genome assemblies by using tools such as Roary [23], Panaroo [24] or PanTA [25]. The latter can construct the graph progressively.

Here we introduce Pasa, the first computational method that makes use of the population information of a species to improve draft assemblies of isolates belonging to the species. Specifically, it uses the pangenome graph built from existing genomes to resolve the assembly graph of a newly sequenced isolate for scaffolding. Pasa builds the former graph by using PanTA [25] on a large set of genomes and using it as the guide to resolve the assembly graphs generated by SPades assembler [26]. We demonstrate the utility of Pasa by applying it to scaffold the draft assemblies of bacterial isolates of species of various levels of genomics diversification, namely, *Klebsilla pneumoniae, Escherichia coli* and *Streptococcus pneumoniae*. In all cases, we show that pangenome graphs are helpful in resolving the assembly graph for better scaffolding quality compared to other multi-reference-based methods.

## 2. MATERIALS AND METHODS

### 2.1. Overview of algorithm

In assembling the target genome sequenced by a short-read technology, an assembler such as Spades [26] and Velvet [27] constructs a *de Bruijn* graph from overlapping reads, and then identifies contigs by finding walks through the *de Bruijn* graph that correspond to continuous sequences. Most genomes of both eukaryotes and prokaryotes organisms contain an abundance of repetitive sequences whose sizes are beyond the length of the reads. In such cases, a walk going into a repetitive sequence has multiple possible paths and thus the corresponding contig cannot be extended unambiguously, leading to the assembly being fragmented into multiple contigs.

Genome scaffolding involves ordering and orientating the contigs in a fragmented assembly and connecting these contigs to improve the assembly’s completeness and contiguity. Pasa (PAngenome-based ScAffolding) achieves this by exploiting the connectivity information obtained from the population genomes of the species. Figure 1 illustrates an overview of the approach of Pasa. In the first step, Pasa runs PanTA [25] on the reference genomes to obtain a pangenome, which is a collection of gene clusters. Pasa then uses the pangenome to build a pangenome graph that captures the structural variant landscape within the population. In this pangenome graph, nodes are clusters of genes and two nodes are connected by an edge if they are adjacent in any genome from the population (Figure 1-A). In scaffolding the assembly of a target genome *T*, Pasa adds genes in contigs in *T* to the population pangenome graph using the add function in PanTA. The pangenome graph essentially establishes the relative positions (that is, distance and orientation) of the contigs in *T*. Pasa then computes the *matching score* for each pair of contigs in *T*. The matching score measures the likelihood of two contigs being adjacent, *i*.*e*., the higher the score between two contigs the more likely they are adjacent in the target genome. In addition, Pasa leverages the information from the assembly graph to obtain a more reliable estimation of the matching scores. Finally, it solves the constrained maximum matching problem to obtain an ordering of the contigs, which is then further refined to obtain the final scaffolds.

**Figure 1.**
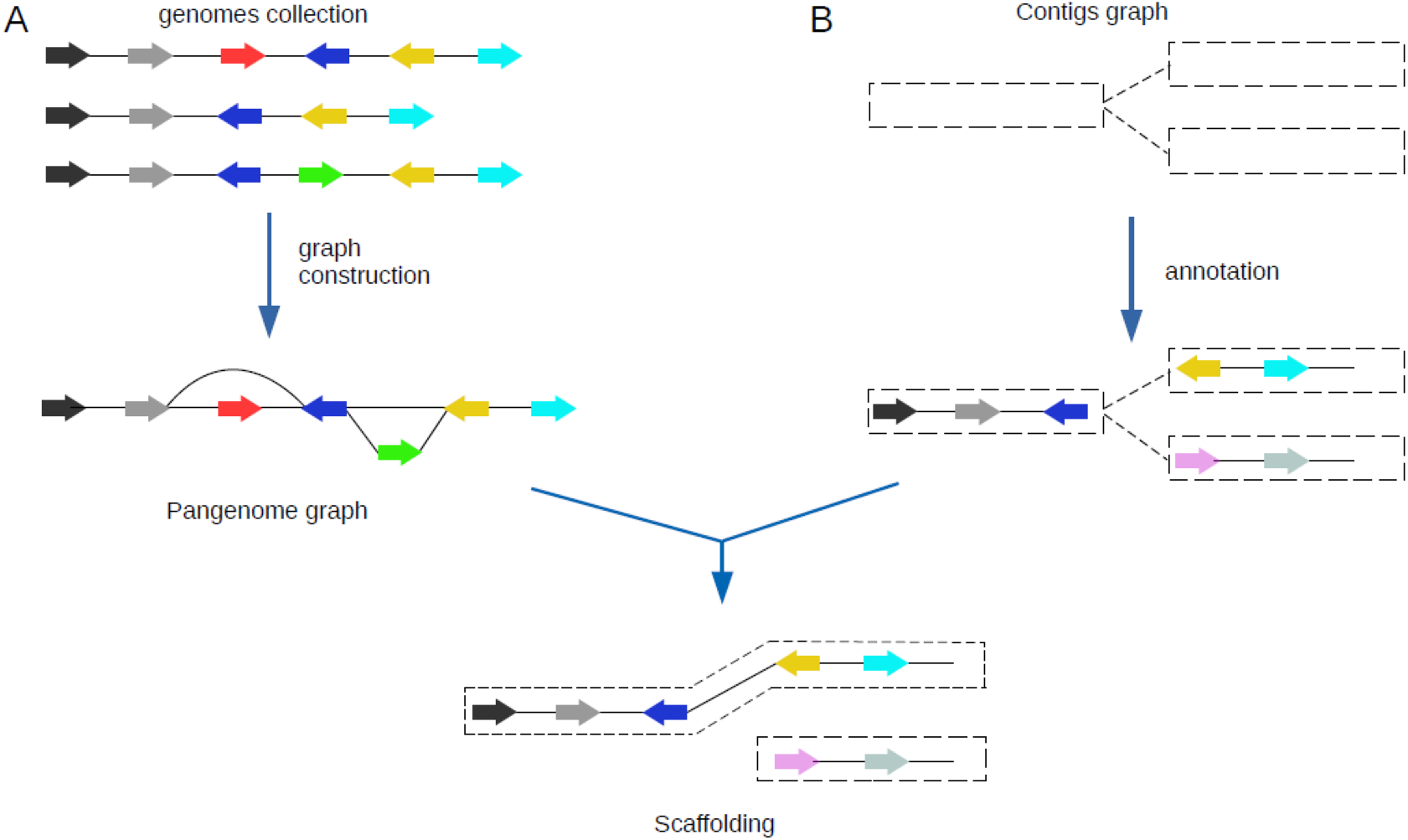
Overview of Pasa algorithm. Genes are represented by arrows and colored by the orthologous groups. (A) Pasa takes the pangenome built from a collection of genomes from a species as input. It then constructs the pangenome graph where nodes represent gene families and edges represent genomic neighborhood. (B) The contig graph represents the possible connections between the contigs in the target genome. Pasa employs information of the gene order of conserved regions in the pangenome graph to resolve the multiple connections in the contig graph.

#### Algorithm 1

Pangraph-based genome assembly

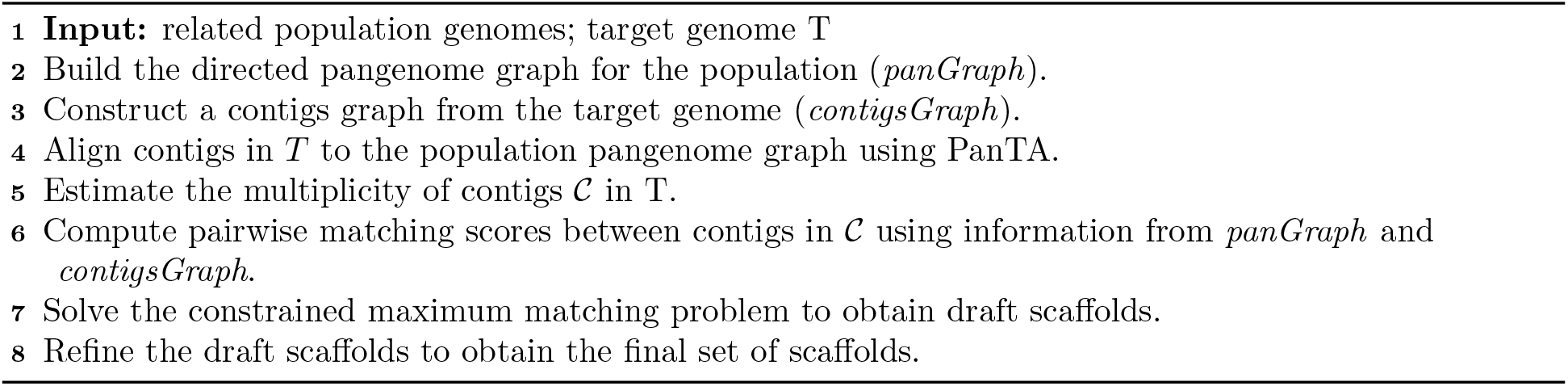

### 2.2. Construction of the pangenome graph

Pasa builds the pangenome graph of the species from the genome assemblies of a collection of isolates. The population genomes are annotated with Prokka [28], which generates a gff file for each genome assembly. Pasa then runs PanTA on the population gff files to obtain gene clusters. Here each cluster represents an orthologous or paralogous group of genes. PanTA groups the genes based on their sequence similarity and outputs a list of genes for each chrosomome or plasmid in the reference genomes. Next, Pasa orients the gene-level genomes obtained by PanTA such that the resulting ordering maximally agrees with each other. Here the agreement is determined based on the number of common pairs of consecutive genes. The orientations of the gene-level genomes are achieved by the following procedure: The algorithm starts with the first genome, and its orientation is arbitrary. Next, Pasa finds an orientation of the second genome that has the largest number of common pairs of consecutive genes with the first genome. Similarly, Pasa finds an orientation of the third genome that shares the largest number of common pairs of consecutive genes with the first two genomes, and the procedure is repeated for the remaining genomes.

Pasa then constructs a directed graph with weighted edges, where nodes represent clusters of genes, two nodes are connected by an edge if they are adjacent in any genome from the population, and the edge weight accounts for the number of times two nodes are adjacent in the oriented genomes. As a result, the edges along the conserved regions of DNA sequences throughout evolution are expected to have substantial weights. Lower weight edges, on the other hand, are likely to correspond to infrequent or under-represented portions of the genome. Pasa keeps edges with high weights and discards those with low weights (less than 20% the number of the reference genomes). It is worth mentioning that Roary [23] and Panaroo [24] also construct a graphical representation of the pangenome, they however only consider a simple undirected unweighted graph to represent the gene arrangements in the population. In contrast, Pasa employs the directed graph with weighted edge because this graphical representation can capture the connectivity information from the population genomes. The edge weights in the pangenome graph are later used to determine the optimal order of the contigs in the target genome.

### 2.3. Construction of contigs overlap graph

Based on the *de Bruijn* assembly graph from SPAdes, Pasa builds a sequence overlap graph of all final contigs, or the *contigs graph* for short throughout the scope of this article to distinguish it from the other graph structures. The SPAdes assembly graph is saved in a FASTG file, namely `assembly_graph.fastg` by default. The sequences in this file are edges from the assembly graph, also known as preliminary contigs before the repeat resolution. The final contigs are then constructed from the consequential repeat resolving step, each comprising a unique path of preliminary contigs when traversing this graph (`contigs.paths`). By combining information from these two files, we construct the graph with contigs as vertices and their *k-mer* overlapping connections as edges.

Formally, Pasa builds a graph 𝒢_*C*_ = {𝒞, 𝒱_𝒞_} where 𝒞 = {*c*_1_, *c*_2_, …, *c*_*n*_} is the set of the final contigs and 𝒱_*𝒞*_ is all possible *k*-overlap edges connecting them. From `assembly_graph.fastg`, we have the graph 𝒢 _*ε*_ = {*E, V*_*E*_} of preliminary contig sequences *E* = {*e*_*i*_}, *i* = 1 … *m* and their connections *V*_*E*_ = {(*e*_*i*_, *e*_*j*_)} for all *e*_*i*_ k-overlap with *e*_*j*_. By investigating the file `contigs.paths`, we know how a final contig is made from a path of the preliminary sequences. For example, if we have 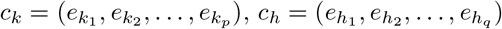 then there is an edge (*c*_*k*_, *c*_*h*_) ∈*V*_*C*_ if and only if 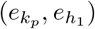 ∈ *V*_*E*_. In the case when intermediate files from SPAdes output are not given but only the final contigs (`contigs.fasta`), Pasa will scan for all possible overlaps between all pairs (*c*_*i*_, *c*_*j*_) ∈ 𝒞. The scanning window for overlapping length is set to the range 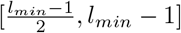 by default, where *l*_*min*_ is the length of the shortest contig amongst 𝒞.

### 2.4. Pangenome graph-based assembly model

#### Alignment of contigs in *T* to the pangenome graph

The draft assembly of the target genome *T* is annotated with prokka and is added into the pangenome graph by the add function in PanTA. When a contig is aligned to the pangenome graph, its orientation is also chosen such that the genes’ direction in the contig and in the pangenome graph agree the most.

#### Multiplicity estimation

To obtain scaffolds as accurately as possible, Pasa must first determine the multiplicity of contigs in the target genome. Pasa uses the length and coverage information to estimate the multiplicity of the contigs. In particular, it utilizes a simplified model from [8]. The median coverage of the five largest contigs is the baseline. The number of copies of the remaining contigs are the ratio of its coverage and the baseline, rounding to the nearest integer.

Intuitively, Pasa creates *k* different copies for each unresolved repeat in the target genome, where *k* is the estimated copy number of the repeat in the complete genome. As some repeats could already be resolved by the NGS assembler, the corresponding synteny blocks in the target contigs will be surrounded by other unique synteny blocks. Pasa uses this “context” information to map repeat instances in contigs to the corresponding repeats in reference genomes. See additional details about the repeat resolution algorithm in the Methods section “Repeat resolution algorithm.”

#### Matching scores

Given a collection of contigs 𝒞 = {*c*_1_, *c*_2_, …, *c*_*n*_}, Pasa assigns a scoring to each pair of contigs in 𝒞. This score function utilizes both population information in the pangenome graph and sample-specific information in the contigs graph. The score indicates how likely the two contigs are joined together, the higher score the more likely they are adjacent in the original genome. The score between two contigs *c*_*i*_ and *c*_*j*_ is defined as

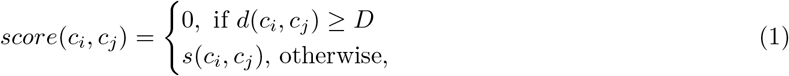

where *d*(*c*_*i*_, *c*_*j*_) is the shortest distance in nucleotides between the two contigs in the contigs graph and *D* is a threshold parameter (*D* = 5000 by default). In addition, *s*(*c*_*i*_, *c*_*j*_) mimics the number of reference genomes that support *c*_*i*_ and *c*_*j*_ to be adjacent. Specifically, *s*(*c*_*i*_, *c*_*j*_) is the average edge weight along a path connecting a terminal gene in *c*_*i*_ and a starting gene in *c*_*j*_ in the pangenome graph, further normalized by the number of genes between the two genes. If there is more than one path between two genes, Pasa takes the shortest one. The score of two contigs is high if they are close to each other in many genomes. Pasa employs multiple genes in the contigs rather than just two end genes to obtain a more accurate score between the two contigs.

#### Constrained maximum matching model

Similar to [17, 18], Pasa splits each contig into a head and a tail node, so that finding an assembly corresponds to a weighted maximum matching in 𝒞. It should be noted that in [17, 18] they only considered the pairwise scores between two contigs, therefore the relationship of more than two contigs is not taken into account. If *score*(*c*_*i*_, *c*_*j*_) and *score*(*c*_*j*_, *c*_*k*_) are high, they will probably join *c*_*i*_ − *c*_*j*_ − *c*_*k*_ regardless of the relationship between *c*_*i*_ and *c*_*k*_. This could be problematic if *c*_*j*_ is a repeat contig. As a result, Pasa exploits long-range dependency between contigs. If *c*_*j*_ is a repeat contig and *s*(*c*_*i*_, *c*_*k*_) ≤ *γ* (*γ* is a parameter), then (*c*_*i*_, *c*_*k*_) is not considered in the matching. Pasa uses a modified version of the greedy algorithm to find the constrained perfect matching (see Algorithm 2).

#### Refinement

There are still certain types of contigs that cannot be considered in the model. These include (1) un-aligned contigs, and (2) short contigs that have no gene. To include these fragments, Pasa uses the contigs graph, which has been constructed from all input contigs. The genome traverses the graph with a certain unknown path. However, since initial scaffolds are now available, Pasa uses these scaffolds to restore small or repetitive fragments. Given a contigs graph and a set of merged scaffolds from the previous step of the algorithm, for each pair of consecutive contigs from these scaffolds, Pasa finds all possible paths connecting them in the contigs graph that do not contain contigs from the scaffold. If there exists such a path, it inserts all the intermediate contigs along this path between the two contigs.

##### Algorithm 2

Constrained scaffolding greedy algorithm

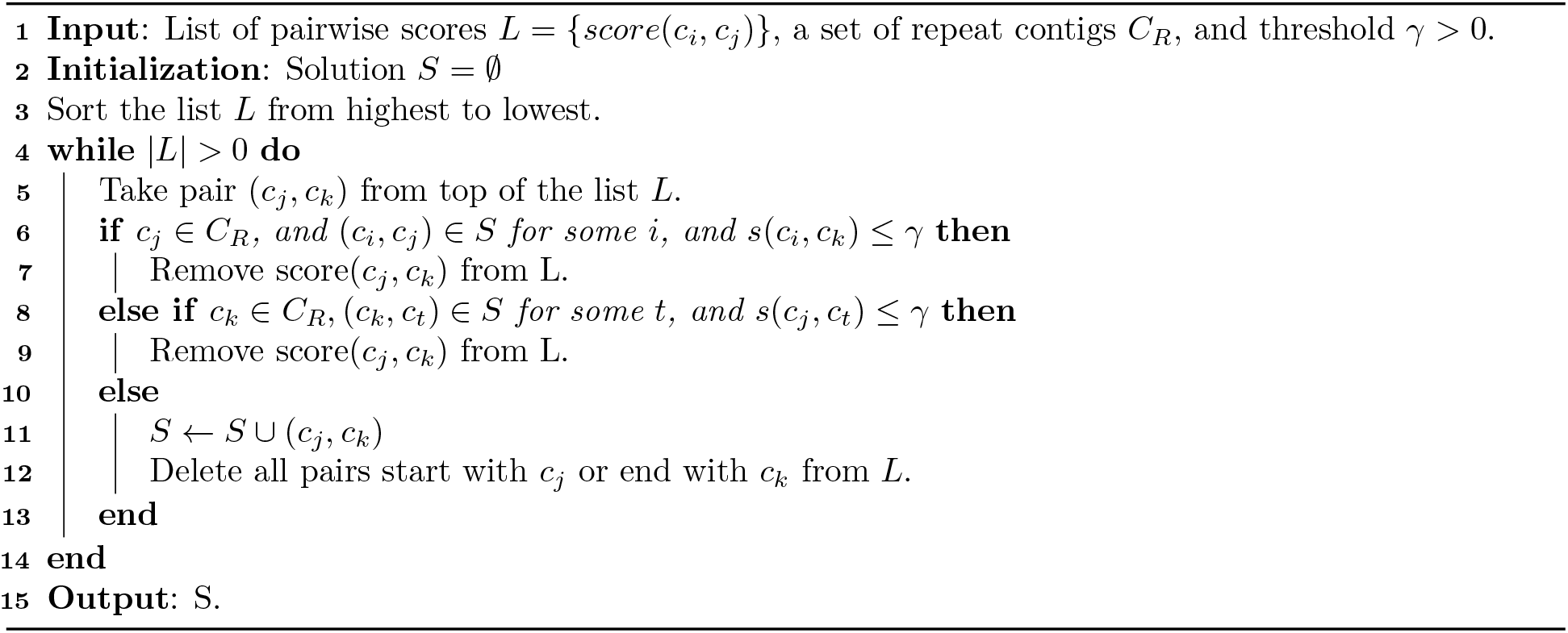

The refinement is further strengthened as follows: For each joined contig from the above refinement step, Pasa considers all single-copy contigs in 𝒞 that made the joined contig, called *backbone contigs*. For two close backbone contigs in the joined contig (i.e. their distance in the joined contig is less than 2 Kbp), the oined contig is split into two at one of the two backbone contigs if their matching score is less than a certain threshold (3.0 by default). This refinement step is omitted in Pasa sensitive mode.

## 3. RESULTS

### 3.1. Evaluation on simulated data

We compared the performance of Pasa to existing state of the art reference-based scaffolders including Multi-CSAR [18] and Ragout (v2) [17]. We excluded MEDUSA [29] from the comparison because it was shown to perform poorly in previous benchmarks [17, 18]. We evaluated Pasa and competing methods on a simulated dataset constructed from the genomes across three different bacterial species, namely *Klebsilla pneumoniae, Escherichia coli*, and *Streptococcus pneumoniae*; these species were chosen to represent differing levels of genomic diversity: conservative, moderate, and divergent, respectively. For each species, we randomly chose 10 complete genomes from the NCBI Reference Sequence Database as the test genomes. The accessions of the genomes are listed in Supplementary Table 1. We also ensured that the test isolates were not included in the set of samples in the reference genomes. The customized Jupyter notebook for downloading and preparing data is included in https://github.com/amromics/pasa.

**Table 1:** Average performance of the evaluated scaffolders on the simulated datasets. For each bacterial species, NGA50, aligned length, and # contig are averaged over the 10 genomes assembled from synthetic sequencing data. All reported numbers are rounded to the nearest integer. The length of the best corrected scaffolds are highlighted in **bold**.

|  | NGA50 (kb) |  |  | aligned length (kb) |  |  | # contigs |  |  |
| --- | --- | --- | --- | --- | --- | --- | --- | --- | --- |
|  | K.p. | E. coli | S.p. | K.p. | E. coli | S.p. | K.p. | E. coli | S.p. |
| Pasa | 1,231 | 408 | 193 | 5,712 | 4,664 | 2,029 | 13 | 27 | 13 |
| Pasa (s) | **1,700** | **558** | **207** | **5,728** | **4,973** | **2,060** | 8 | 15 | 8 |
| Ragout | 257 | 188 | 158 | 4,222 | 4,061 | 1,968 | 14 | 12 | 7 |
| Multi-CSAR | 1,190 | 408 | 172 | 5,642 | 4,530 | 2,037 | **5** | **8** | **4** |

For each test genome, we simulated Illumina sequencing reads using ART (v2.5.8) [30] with the following configuration: paired-end sequencing, read length of 100bp, fold coverage 70 and mean fragment size of 400bp. We ran SPades assembler (v3.13.0) [26] on the simulated sequencing data to construct the draft assembly for the genome. We then applied the competing methods to scaffold the draft assembly using the nformation from the reference genomes of the species.

We evaluated the scaffolders using common metrics including NGA50 statistics, the aligned length, the number of resulting contigs and the number of misassemblies. While the NG50 metrics is the length (in Kbp) for which the collection of all contigs of that length or longer covers at least 50% of the genome, the NGA50 metric is similar, but uses aligned blocks instead of contigs for the calculation. Total aligned length is the total number of aligned bases in the scaffolds. This value is usually smaller than a value of total length because some contigs may be unaligned or partially unaligned to the reference. In principle, NGA50, the aligned length, and the contig number are metrics to access the contiguity of scaffolds whereas the number of misassemblies measures the accuracy of the scaffolding. These metrics were calculated using QUAST v5.0.2) [31].

Table 3.1 and Figure 2 report the average performance of the scaffolding methods on the test genomes for each species. We found that Pasa (s) in sensitive mode achieved the most complete assemblies, as measured by NGA50 and the aligned length metrics. Pasa and Multi-CSAR showed similar performance in terms of contiguity of scaffolds, which both outperformed Ragout (Table 3.1). The NGA50 metrics of Ragout were consistently low across all test genomes. Ragout’s usage of synteny blocks to bridge the contigs may be one of the causes. Because these synteny blocks are frequently short, only a small number of contigs are linked n Ragout.

**Figure 2.**
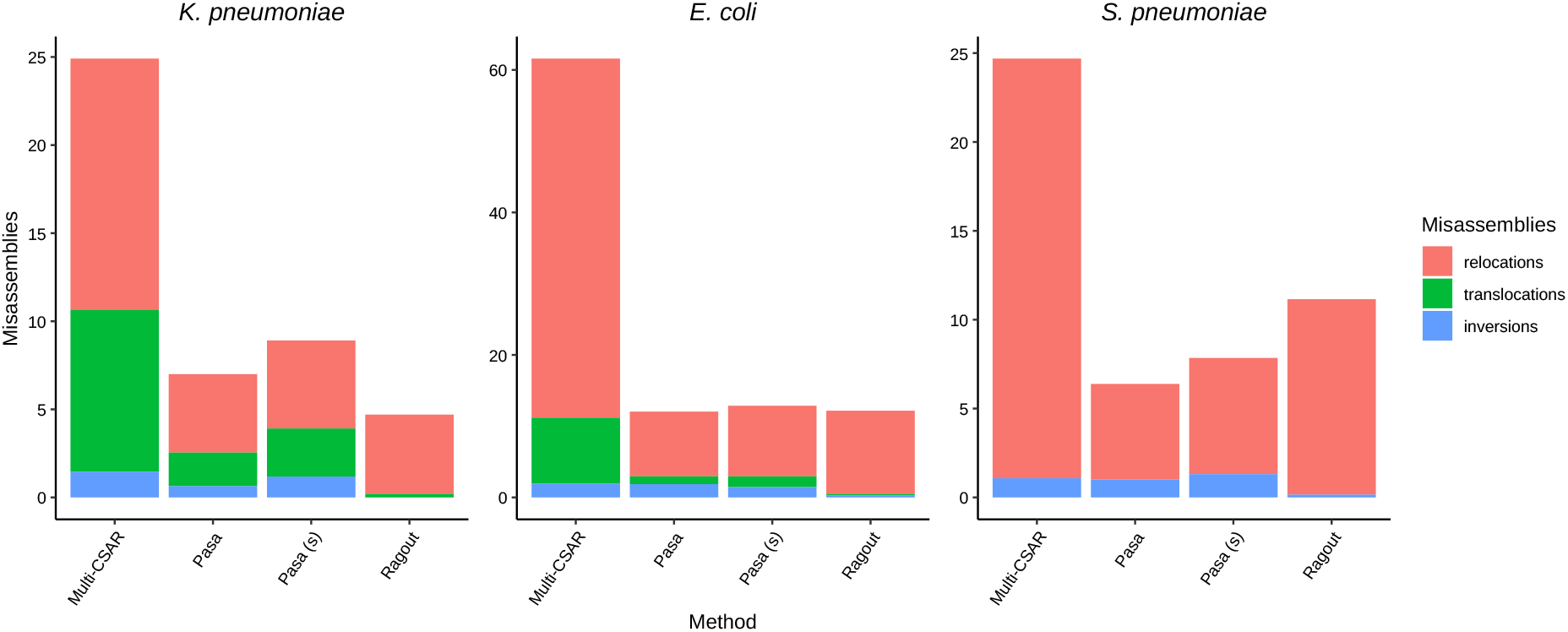
Average number of misassemblies (relocations, translocations, inversions) of Pasa and competing methods on ten datasets across three different strains. A relocation is a misassembly event (breakpoint) where the left flanking sequence and the right flanking sequence align overlap or away from each other on the same chromosome of the reference genome. A translocation is a misassembly event (breakpoint) where the flanking sequences align on different chromosomes. Inversion corresponds to a breakpoint where the flanking sequences align on opposite strands of the same chromosome.

Multi-CSAR is compatible with both versions of Pasa in terms of contiguity metrics, it however produced the largest number of misassemblies, generally 4-6 times higher than Pasa and Ragout (Figure 2). The misassembly rates of Pasa and Ragout were very low since both methods incorporate the connectivity information in the assembly graph into the pipeline. Specifically, Ragout made fewer errors than Pasa in the *K. pneumoniae* dataset but was more erroneous than both versions of Pasa in the other bacterial species. Figure 2 shows that relocations are the most common source of mistakes, which is expected given that all techniques employed reference genomes to bridge the contigs, and those reference genomes may have different gene orders than the target genome, resulting in the incorrect order of contigs in the scaffolds. Both Multi-CSAR and Pasa made few translocations and inversions misassemblies, and the number of errors is low (typically *<* 3).

As expected, Pasa in its conservative mode is less likely to produce a complete assembly than its sensitive mode but it carries a low risk of misassembly and is appropriate for contexts where assembly accuracy is important. In sensitive mode, the refinement is less restricted. This mode is most likely to complete the assembly but carries a slightly greater risk of error. It is suited to cases where completeness is more important than accuracy. In summary, Pasa and Ragout performed similarly in terms of misassemblies, whereas Pasa outperformed Ragout in terms of contiguity (up to 6 folds in *K. pneumoniae* isolates). Despite the fact that Multi-CSAR produced fewer contigs than Pasa and performed similarly to our method in terms of contiguity, the misassembly rates were substantially greater, highlighting the difficulties of resolving repeats using the reference alone.

### 3.2. Evaluation on real data

Next we compared Pasa to the competing methods on real data where the genomes were sequenced using Illumina technology. We obtained sequencing data for three isolates belonging to three species: *Staphylococcus aureus, Klebsilla pneumoniae* and *Salmonella enterica*. For each isolate, we ran the competing pipelines on the shortread sequencing data and benchmarked the results against the complete genome. The complete genome and the sequencing data for the *S. aureus* isolate were obtained from the Genome Assembly Gold-Standard Evaluation (GAGE) benchmark study [32]. The *S. aureus* dataset in the GAGE benchmark was previously sequenced and finished using conventional Sanger technology, and later resequenced using Illumina technology. For the other two species, we selected the complete genomes of two isolates from the RefSeq database with Accession number GCF 003030145.1 and GCF 000439415.1, respectively. The sequencing data of these two isolates were obtained from the SRA archive database (Run accessions: SRR9042857, SRR9043663)

We used NGA50 metrics and the number of misassemblies as the representative metrics for contiguity and accuracy. Figure 3 summarizes the performance of the scaffolding tools on real datasets. Specifically, for the *S. aureus* dataset from the gold-standard database, we found that Pasa made fewer incorrect joins than all other scaffolders while achieving the best scaffolding results in NGA50 metrics (Figure 3-left). Note that this dataset had a low NGA50 metric due to the old sequencing technology (Illumina Genome Analyzer II, average read length: 101 bp, reads coverage: 45). For the *K. pneumoniae* dataset, both Pasa and Pasa (s) were significantly more accurate than Multi-CSAR; they produced genome assemblies with 10-fold and 2-fold fewer missassemblies, respectively while still exhibiting slightly lower NGA50 metrics (Figure 3-center). It can be seen that the NGA50 metrics and the number of misassemblies of Pasa and Ragout were consistent with the report of *K. pneumoniae* in the simulation study. This indicates that our simulation strategy captures real data well. For the *S. enterica* dataset, Pasa (s) outperformed the competing methods in terms of NGA50 metrics. The NGA50 metrics of Pasa (s) were around 2800Kp, which were approximately four times higher than Multi-CSAR (≈ 700Kp) and seven times higher than Ragout (≈ 400Kp). In addition, the numbers of misassemblies of Pasa and Pasa (s) were the same, and were slightly higher than Ragout but much lower than Multi-CSAR (Figure 3-right). In summary, the results of real datasets were consistent with the simulated datasets. Multi-CSAR generally produced the largest number of misassemblies and Ragout had the worst performance in the NGA50 metrics. In contrast, Pasa generally produced more complete assemblies than the competing methods while maintaining a low error rate.

**Figure 3.**
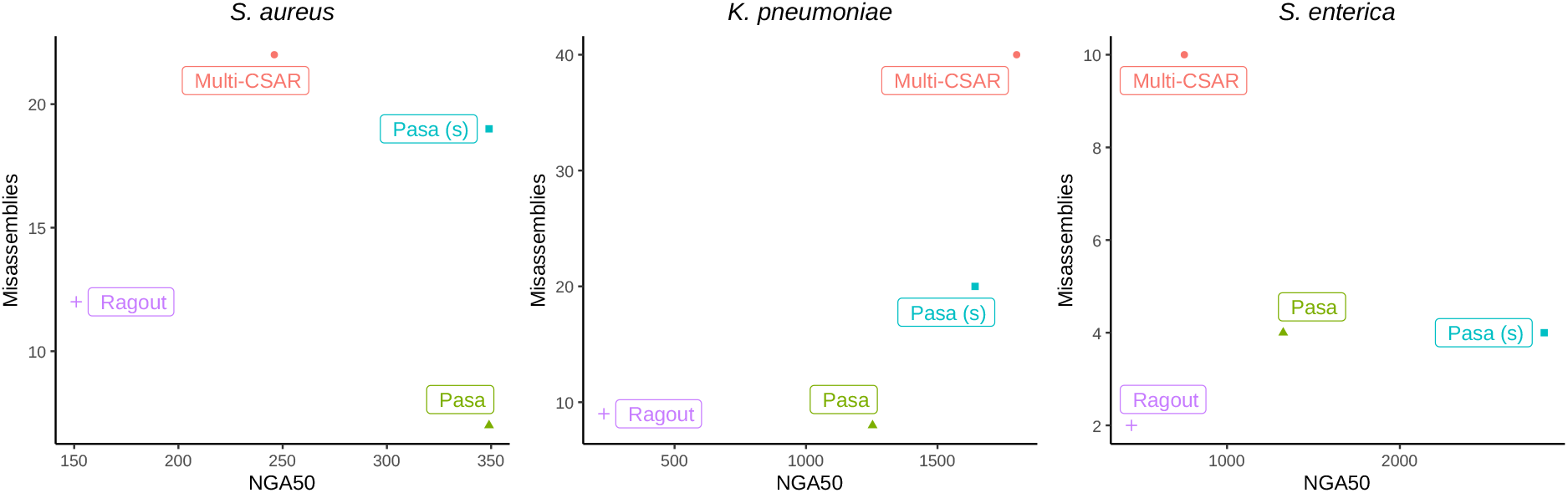
Performance of scaffolders on real datasets. Scatter plots shows the NGA50 metrics and the number of misassemblies made by Pasa and competing methods on *S. aureus* (left), *K. pneumoniae* (center), and *S. enterica* (right).

### 3.3. Scaffolding using the pangenome of a related species

When reference pangenome genomes are unavailable, a closely related species can be used as a substitute. Scaffolding using a related reference pangenome is crucial in practice since reference genomes for some species, particularly uncommon and novel bacterial species, are not always available. To show our scaffolder’s ability to use related reference pangenomes, we applied Pasa to scaffold the genome assembly of a *K. quasipneumoniae* isolate using the pangenome constructed from a population of *K. pneumoniae* species as the reference. We compared the results with Multi-CSAR and Ragout. The results are shown in Figure 4. Despite using a different reference, Pasa obtained reasonable results in terms of contiguity and accuracy. In particular, the NGA50 metric of Pasa (sensitive) was twice that of Multi-CSAR and 6-7 times higher than that of Ragout. Ragout had the lowest number of misassemblies, followed by Pasa and Pasa (sensitive). When the target genome was assembled using the same reference as the target, the NGA50 metrics of Pasa and Multi-CSAR increased significantly (Figure 4-right). Accordingly the NGA50 metrics of Pasa and its sensitive version were still better than that of Multi-CSAR and exceeded that of Ragout. Pasa had slightly more misassemblies than Ragout but less than Multi-CSAR. Overall, this shows the practical usage of our scaffolder, even if references are not available.

**Figure 4.**
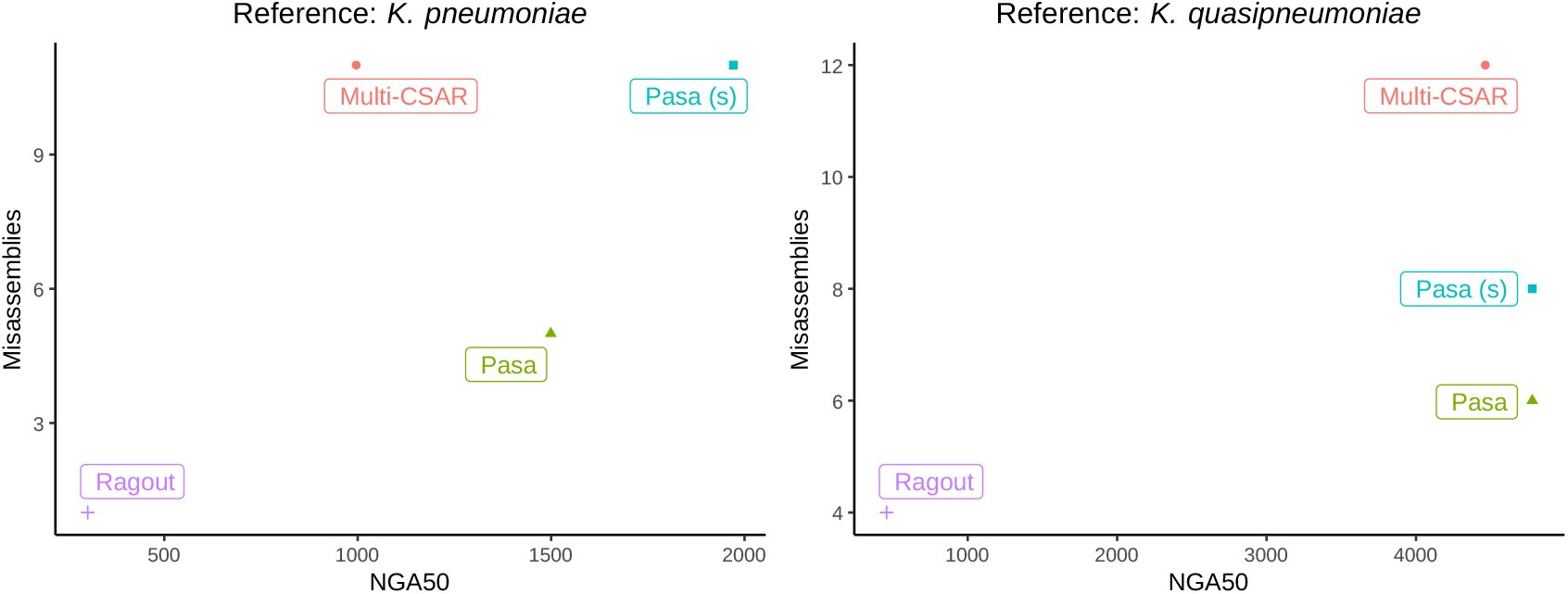
Performance of scaffolders using the related and exact reference. Scatter plots shows the NGA50 metrics and number of misassemblies made by Pasa and competing methods on *K. pneumoniae* (close) and *K. quasipneumoniae* (exact).

### 3.4. Time and memory usage

We demonstrate the scalability of Pasa to large data sets. Table 2 gives the average CPU times (in minutes) and peak memory usage (in GB) of the scaffolders on the synthetic data sets from the three bacterial species used in this study. Since the running time and memory usage of Pasa and its sensitive mode were similar, we only reported the performance of Pasa. It can be shown that Pasa completed a genome in around 2 minutes, which was 8 times faster than the second fastest method, Multi-CSAR. Note that the running time of Pasa did not include the running times of the pangenome construction steps because the reference pangraph only needs to be built once and can then be applied to datasets of the same bacterial species. Nevertheless, the total running time of Pasa and the pangenome construction steps (Pasa+Pangraph) was around 4 minutes, which was approximately four times faster than Multi-CSAR. Ragout was the slowest, taking around 2 hours to finish a genome. Table 2 (right) shows peak memory usage for the same simulated data sets. Multi-CSAR required the least amount of memory (*<* 0.2 GB), followed by Pasa (0.6 − 0.8 GB) and Pasa with the pangenome construction steps (around 1.3 GB). Ragout used substantially more memory (7 − 9 GB) than the other methods.

**Table 2:** Comparison of running times and peak memory usage on 30 synthetic data sets. Average running times (in minutes) and memory usage (in GB) of Pasa, Multi-CSAR, and Ragout are reported on the synthetic data sets used in this study. *Running times exclude the construction of the pangenome graph step as it is done only once and reused for the later genome. The last row (Pasa+Pangraph) reports the total running times and peak memory usage of Pasa and the pangenome graph construction step.

|  | Runtime (minutes) |  |  | Memory (Gb) |  |  |
| --- | --- | --- | --- | --- | --- | --- |
|  | K.p. | E. coli | S.p. | K.p. | E. coli | S.p. |
| Pasa | 2.08 | 1.93 | 1.85 | 0.73 | 0.63 | 0.60 |
| Ragout | 121.23 | 117.15 | 111.54 | 8.66 | 7.38 | 7.28 |
| Multi-CSAR | 17.06 | 16.63 | 15.42 | 0.19 | 0.17 | 0.16 |
| Pasa + Pangenome | 4.38 | 4.12 | 3.98 | 1.35 | 1.31 | 1.26 |

## 4. DISCUSSION

Obtaining the complete genome assemblies is important in microbial genomics, especially in the context of antimicrobial resistance research and surveillance. Complete genomes provide crucial information of gene positions in the chromosomes and plasmids to elucidate the development and transmissions of drug resistance. Pasa offers an efficient way to readily improve the contiguity and completeness of the large number of bacterial genomes that are sequenced by the affordable and high-throughput Illumina sequencing technology. Instead of relying on extra laboratory experiments that are time consuming and costly, Pasa utilizes the connectivity information of the existing population genomes to resolve the assembly graph of the short-read assembly. Pasa is also fast and efficient, making it suitable to scaffold the large number of short read assemblies available.

The use of population information for the scaffolding problem has been exploited in several reference-based scaffolder methods such as Multi-CSAR and Ragout. When compared with the two methods, the benefits of using Pasa are twofold: first, it produces a more complete genome with fewer errors; second, it has the lowest running time. The key difference between Pasa and scaffolders such as Multi-CSAR and Ragout is that Pasa employs a compact graphical representation of the pangenome from the population genomes. In addition, Pasa solves the constrained maximum matching problem, which allows modeling longer range dependencies between contigs in the target genome. The method may further be improved by developing an exact algorithm for the constrained maximum matching problem. In the future, it will be interesting to investigate the performance quality of our tool Pasa using different assemblers as well.

Prokaryotic genomes are known for enormous intraspecific variability owing to great variation events such as horizontal gene transfers, differential gene losses and gene duplication [33]. Pangenome analysis was introduced as a methodology to capture the diversity of bacterial genomes [34] and has been a dispensable tool in microbial genomics studies [35] to generate biological insight such as understanding the evolution of bacterial species [36, 37], variant detection [38] and studying‘’ antiobiotic resistance [39]. In this work, we extend the utility of the pangenome by introducing a novel method to exploring the species information to scaffold and improve the quality of new genome assemblies.

The construction of the pan-genome graph allowed us to learn the gene order information in a population. The conservation of gene order can thus be used to help order contigs along a chromosome by inferring their placement based on the location within the pangenome graph of the genes found in the contigs. The target genome however diverges from the population, we therefore also explored the sample-specific information by means of the assembly graph constructed from the target genome. Exploiting the information from both sample-specific information and population resulted in a highly efficient scaffolding method, Pasa. We demonstrated that Pasa could improve the contiguity and accuracy of the genome scaffolding. We observed consistent results on both real and simulated datasets for various bacterial species. Furthermore, we demonstrated that Pasa does not require the reference genomes to be the same species as the target genome by utilizing related genomes as the reference. This is especially useful when the reference genomes are unavailable.

The vast majority of existing microbial genome assemblies are produced by short-read sequencing technology; many more genomes are being sequenced in research laboratories, medical organizations and public health agencies. Pasa is developed to meet the need of efficient bioinformatics methods to make sense of the data. It was demonstrated to be able to improve the contiguity and accuracy of the genome scaffolding by using the information from both the pangenome graph and the contig graph. We observed consistent results on both real and simulated datasets for various bacterial strains. Pasa offers an accurate and efficient method to scaffold and improve the completeness and continuity of these genomes, without the need for extra laboratory experiments.

## SOFTWARE AVAILABILITY

The Pasa software is available at GitHub (https://github.com/amromics/pasa) under the open-source MIT license. The Pasa repository also includes all code necessary to reproduce the results of this manuscript.

## ACKNOWLEDGEMENTS

H. Do would like to thank the Vietnam Institute for Advanced Study in Mathematics (VIASM) for the hospitality and for the excellent working conditions.

