## Supplemental Table 1 for "Pasa: Leverage population pangenome graph to scaffold prokaryote genome assemblies"

**Accessions of the isolates used in the study**

**Klebsilla pneumoniae**

https://www.ebi.ac.uk/ena/browser/view/GCA_001705385.1

https://www.ebi.ac.uk/ena/browser/view/GCA_021390015.1

https://www.ebi.ac.uk/ena/browser/view/GCA_021249265.1

https://www.ebi.ac.uk/ena/browser/view/GCA_021166135.1

https://www.ebi.ac.uk/ena/browser/view/GCA_021166095.1

https://www.ebi.ac.uk/ena/browser/view/GCA_004322955.1

https://www.ebi.ac.uk/ena/browser/view/GCA_001521895.1

https://www.ebi.ac.uk/ena/browser/view/GCA_001456135.1

https://www.ebi.ac.uk/ena/browser/view/GCA_001307175.1

https://www.ebi.ac.uk/ena/browser/view/GCA_000814805.1

**Escherichia coli**

https://www.ebi.ac.uk/ena/browser/view/GCA_900174635.1

https://www.ebi.ac.uk/ena/browser/view/GCA_023658305.1

https://www.ebi.ac.uk/ena/browser/view/GCA_022918835.1

https://www.ebi.ac.uk/ena/browser/view/GCA_020526805.1

https://www.ebi.ac.uk/ena/browser/view/GCA_018075345.1

https://www.ebi.ac.uk/ena/browser/view/GCA_016864175.1

https://www.ebi.ac.uk/ena/browser/view/GCA_015571535.1

https://www.ebi.ac.uk/ena/browser/view/GCA_014295255.1

https://www.ebi.ac.uk/ena/browser/view/GCA_014169015.1

https://www.ebi.ac.uk/ena/browser/view/GCA_013167255.1

**Streptococcus pneumoniae**

https://www.ebi.ac.uk/ena/browser/view/GCA_022318405.1

https://www.ebi.ac.uk/ena/browser/view/GCA_022075545.1

https://www.ebi.ac.uk/ena/browser/view/GCA_022070425.1

https://www.ebi.ac.uk/ena/browser/view/GCA_022069545.1

https://www.ebi.ac.uk/ena/browser/view/GCA_022069445.1

https://www.ebi.ac.uk/ena/browser/view/GCA_022068405.1

https://www.ebi.ac.uk/ena/browser/view/GCA_019456615.1

https://www.ebi.ac.uk/ena/browser/view/GCA_008253725.1

https://www.ebi.ac.uk/ena/browser/view/GCA_003966525.1

https://www.ebi.ac.uk/ena/browser/view/GCA_000251085.1
